# Variability vs Phenotype: multimodal analysis of Dravet Syndrome Brain Organoids powered by Deep Learning

**DOI:** 10.1101/2025.01.28.635291

**Authors:** Isabel Turpin, Adriana Modrego, Andrea Martí-Sarrias, Anna-Christina Haeb, Laura García-González, Jordi Soriano, Núria Ruiz, Irene Peñuelas-Haro, Elisa Espinet, Daniel Tornero, Oscar Lao, Sandra Acosta

## Abstract

Brain organoids (BO) have risen as a reliable model for neurodelopmental disorders (ND), reproducing human brain development milestones. However, their significant intra- and inter-organoid variability compromises their use in advanced tasks such as drug testing. Overcoming experimental variability is crucial for models prone to variation, like unguided BO. BO modelling in Dravet Syndrome, a late-onset epileptic ND, represents a great challenge since BO variability accumulates with time, when phenotype shows in vitro. Leveraging deep learning, we developed ImPheNet, a predictive tool grounded in BO live imaging datasets. ImPheNet accurately classified phenotypes and assessed drug toxicity in BO derived from DS, revealing differences between genotypes and upon antiseizure drug exposure. These results are supported by transcriptomic and functional data, revealing an excitatory-inhibitory imbalance during the maturation of DS organoids. Altogether, our DL-predictive live imaging strategy, ImPheNet, emerges as a powerful tool enhancing BO research and advancing ND treatments.

## INTRODUCTION

The use of human brain organoids has become a widespread tool to model neurodevelopmental disorders *in vitro*, due to their potential to reproduce in a simpler, more accesible and inferable manner the physiological functions inherent of the human brain, both in healthy and pathological contexts. These advantages stem from the presence of a reliable maturing neurodevelopmental 3D cellular structures growing in suspension, directly derived from human donors and/or patient-specific cells ^1–6^. Despite being described more than a decade ago, organoids widespread applicability has been hindered by two primary challenges: their predominantly immature phenotype and their intrinsic variability ^2,7^. The inherent variability in brain organoids manifests at multiple levels: among organoids produced in the same batch; between experimental replicates (batches); and even more so, between different pluripotent stem cell (PSC) lines. Efforts to strike a balance between diversity and reproducibility are essential for maximizing the potential of brain organoids as reliable and scalable models for studying human brain development and disease. With this purpose, various protocols for the generation of brain organoids have been proposed, broadly categorized into two approaches: guided and unguided. Guided organoids are more reproducible but show reduced cellular diversity ^3,4^. Unguided brain organoids are characterized by high cellular diversity, better recapitulation of the layered cortical plate, but strong variability between organoids even intraexperimentally ^1,8–10^. The successful identification of phenotypic traits in brain organoids strongly relies on selecting the appropriate protocol, carefully balancing its advantages and disadvantages. In unguided protocols, two key stages contribute to the observed variability: the early neural induction and the establishment of the neuronal networks ^11–13^. During early developmental stages, variability can lead to the stochastic generation of multiple regionalized areas and suboptimal cerebral organoids ^11,14^, enriched with non-neural cell types and aberrant morphological traits, producing suboptimal brain organoids. The origin and relevance of variability arising at later stages, when the conformation and the extension of the neuronal circuitry and its functional connectivity is being established, remain largely unexplored. It is associated to the generation of multiple neuronal subtypes, driven by cell-autonomous and non-cell autonomous mechanisms and the heterochronic maturation of the multiple neurogenic rosettes ^3,11,15^. Moreover, it remains unknown to what extent these inherent differences affect them at maturer stages, ballasting the use of brain organoids in late-onset disease modeling. By addressing these challenges, brain organoids can be harnessed to enhance our understanding of complex neurological processes and pave the way for novel therapeutic approaches.

Developmental and epileptic encephalopathies (DEE) encompass a group of disorders characterized by frequent intractable seizures, brain developmental and functional defects, cognitive deficits, and, in some cases, premature mortality ^16^. Dravet Syndrome (DS), the most common DEE, is associated with heterozygous mutations in SCN1A gene, which encodes one subunit of the Nav1.1 sodium channel found primarily in inhibitory neurons ^17–19^. The epileptic phenotype in DS is caused by an imbalance in excitatory-inhibitory circuitry, as a result of decreased sodium current in inhibitory neurons ^20,21^. Other severe DEE have been successfully modeled using brain organoids, such as Rett Syndrome, where pathogenic epileptogenic currents were observed compared to isogenic cell lines ^22^. In this context, the complexity and uniqueness of cerebral organoids analyzed through conventional molecular and histological techniques faces significant challenges in identifying non-obvious phenotypes. Thus, leveraging Artificial Intelligence (AI)-based analyses of whole organoids emerge as a potential solution to overcome the intrinsic variability associated with human brain organoids during phenotyping approaches. AI has appeared as a transformative force in various fields, and its application in life sciences has revolutionized the way we approach research, diagnosis, and treatment ^23,24^. Machine learning and deep learning (DL) algorithms have shown tremendous potential in handling vast amounts of complex biological data and extracting valuable insights. In life sciences, AI-driven approaches are being used to analyze genomics, transcriptomics, proteomics, and other high-throughput data, enabling researchers to identify patterns, biomarkers, and novel therapeutic targets ^25^. Additionally, AI has proven its efficacy in predicting drug interactions, designing personalized treatment plans, and even discovering new drug candidates ^26^. By merging advanced computational methods with biological knowledge, AI is accelerating scientific discovery and paving the way for more precise and personalized approaches to address various health challenges.

In this study, we model DS using brain organoids and determine an underlying epilepsy-associated phenotype in their abnormal neuronal connectivity. Subsequently, we apply DL algorithms to leverage the analysis potential of brain organoids to define the pathological phenotype using live-microscopy images. The multimodal analysis allows to discriminate between inherent brain organoid variability and genotype-associated phenotype. Using this approach, we determine that the phenotype rises above the intra- and inter-organoid variability and predict the potential toxic impact of anti-seizure drugs in DS. Altogether, we define the DS phenotype in brain organoids and we propose a robust methodology based on live-imaging and DL to boost the applications of brain organoids in translational and biomedicine and therapeutic discovery.

## RESULTS

### Temporal and regional variability in human brain organoids

Unguided brain organoids differentiation follows the “inside-out” pattern of cortical differentiation, similarly to human cerebral cortex development where neurons are generated in an inner region and then migrate outward to form distinct layers. This process is heterochronic in brain organoids and maturation rates differ amongst the regional structures in an uncontrolled manner. Here, we aim to monitor the maturation rates using live imaging in brain organoids. To follow maturation changes, human embryonic stem cells (hESC) H9-derived brain organoids were stained at multiple stages (day 35-progenitor expansion and neurogenesis onset, day 60-neurogenesis and day 90-maturing neuronal network and astrogenesis) with Cellbrite, a DiO-based neuronal tracer with reduced toxicity (Figure 1A). This staining resolved organoid surface morphology and neural cytoarchitecture more accurately than brightfield (Figure 1 A-D, Supplementary Figure 1A-C and Supplementary video 1A). Moreover, Cellbrite highlights the surface topography variability and the shape morphological differences among multiple organoids, and, remarkably amongst the multiple regional levels in the same organoid (Figure 1C,D). Limited Cellbrite staining was detected in young organoids at day 35 (Figure 1A), enriched in neural progenitors, supporting its use for evaluating the neural circuitry in mature organoids.

**Figure 1.**
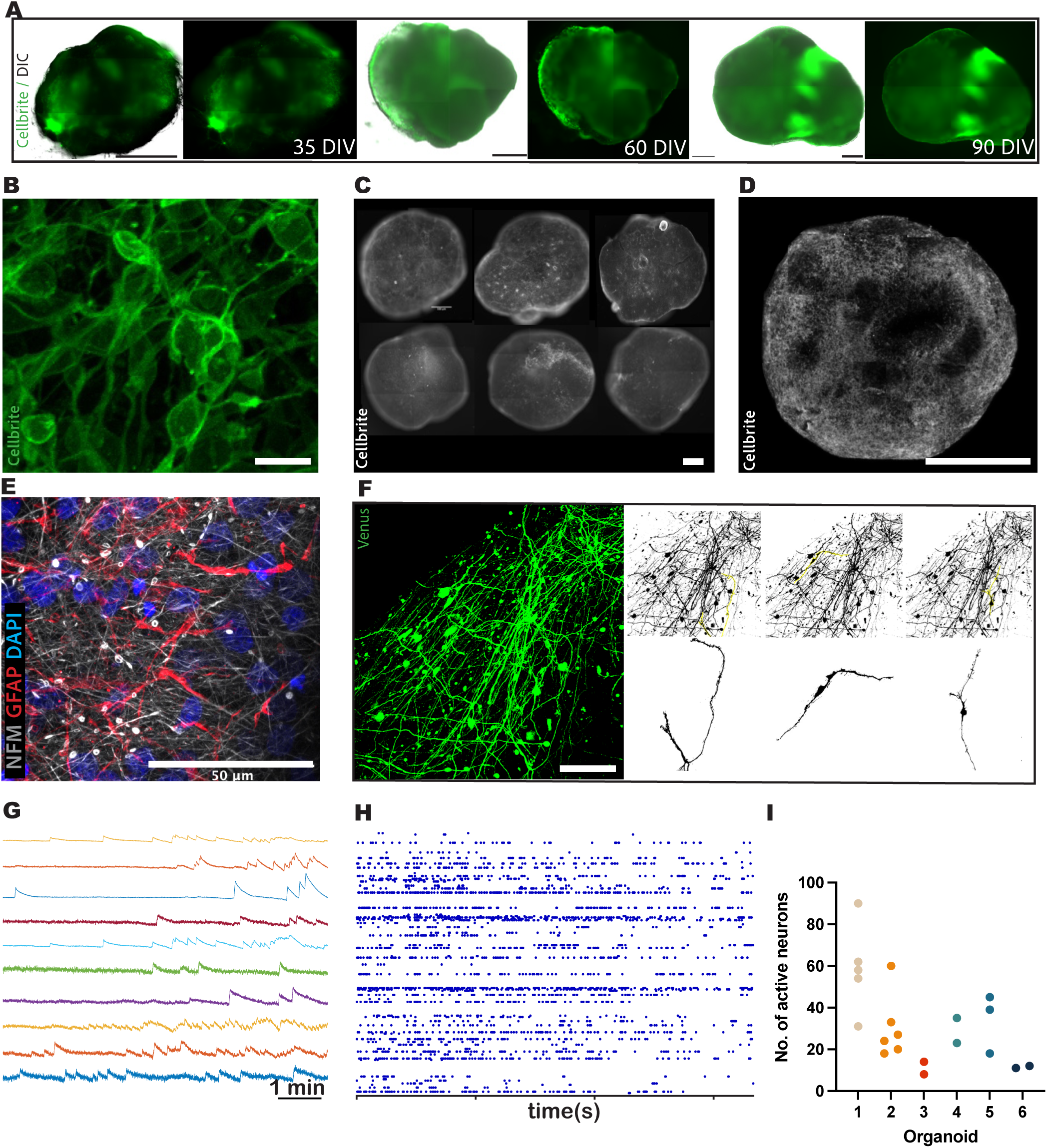
Morphological and functional reconstruction of intra-organoid variability. A. Merged brightfield and Cellbrite fluorescent images from H9 brain organoids at days 35, 60 and 90 of differentiation. Images obtained using Axio Observer Z1 Apotome inverted fluorescent microscope. Scale bar: 500 µm. DIV: days *in vitro*. B. Live-imaging 90-day-old brain organoid stained with Cellbrite. Scale bar: 20 µm. C. Inter-organoid variability of H9 brain organoids at day 90 of differentiation stained with Cellbrite. Scale bar: 250 µm. D. Whole-organoid maximal projection reconstruction of a high-magnification image of a fixed 90-day-old brain organoid stained with Cellbrite. Scale bar: 700 µm. E. Immunofluorescence (IF) staining of NFM and GFAP markers showing mature neurons and astrocytes respectively. Image deconvolution was performed with Airyscan2 inverted confocal microscope-associated software. Scale bar: 50 µm. F. Live-imaging (left) and single neuron segmentation (right) of low-dosage infection with a lentiviral Venus reporter on a day 90 brain, showing heterogeneity in cell morphology and distribution accross the surface. G. Calcium signaling activity traces of individual Syn1+ neurons from a 90-day-old brain organoid infected with a GCaMP6s AAV. H. Raster plot showing firing activity of individual Syn1^+^ neurons over 5 minutes. I. Dotplot indicating the number of active (GCaMP6s) neurons per recorded region of individual organoids.

Co-staining for Cellbrite and markers of neuronal and glial identity indicated the presence of Cellbrite positive staining in both immature neurons and astrocytes in 90 days old brain organoids (Supplementary Figure 1C-E). Moreover, the neural cytoarchitecture at day 90 brain organoids analyzed 1 week after Cellbrite staining was conserved, as the intermingled neuronal (MAP2, Tuj1, NFM) and astrocyte (GFAP) populations showed, (Figure 1E and Supplementary Figure 1D-G), altogether supporting the use of Cellbrite as a suitable live imaging staining for longitudinal neural network studies. At the cellular level, the maturation stage at day 90 of differentiation becomes evident in H9 brain organoids, as they exhibit a well-developed neuronal network with axonal terminations that connect multiple regions within the organoid (Figure 1B-F and Supplementary Figure 1B) highlighting the morphological subregional variability.

Transcriptomic analysis at multiple differentiation stages (day 21, 35 and 90) confirmed that, by day 90, organoids acquire an advanced maturing stage and differ significantly from day 21 and 35 brain organoids, which show an immature neural identity (Supplementary Figure 2A-D). Genes involved in neuronal differentiation, maturation, and connectivity signatures changed over time. Early progenitor signature genes SOX2, NES and ASPM were expressed more strongly at earlier stages, whilst genes indicating mature neuronal signature, such as SYN1, MAP2, DLG4 (PSD-95) and NEUROD6 showed an increasing trend of gene expression from day 21 to day 90. Similarly, neuronal cortical layer markers of deep (TBR1 and BCL11B) and upper layers (POU3F2 and SATB2) showed a sustained increase in expression, alongside markers for late-stage astrocytes (i.e. AQP4 and GFAP). Still, the elevated expression at day 90 of immature neural markers (TUBB3 for early born immature neurons, FABP7 for RG and EOMES for intermediate progenitors) suggests the remarkable heterochronicity in the entire process of organoid differentiation.

While neurogenesis heterochronicity in unguided brain organoids has been observed previously ^11^, it remains unknown how it impacts the connectivity and neuronal function. Thus, we performed detailed analysis of the nature of the intra-organoid connectivity by labelling organoids with low dosage Venus-lentiviral reporter. These results revealed a non-homogeneous distribution of the morphologically distinct pyramidal and bipolar interneurons and their connections in the whole organoid (Figure 1F). The axo-dendritic network appeared to be more densely distributed in certain regions, forming valley-like paths between adjacent vesicle-like structures within the organoid (Figure 1F, Supplementary Figure 1H and Supplementary Video 1B). This distinctive distribution of axonal circuitry significantly contributes to the regional variability observed in brain organoid morphology.

Next, we analyzed neuronal activity within the organoids through the monitorization of intracellular calcium levels. To this purpose, we infected 90 days old organoids (n=6) with AAV7m8 viral vector for the expression of the genetically encoded calcium indicator GCaMP6s under the control of Syn1 promoter. Upon analyzing multiple 10 minutes-long calcium recordings taken at different fields of view within each organoid, it became evident that neuronal activity was mature and rich (Figure 1G-H) and neurons generated complex networks, with synchronized depolarization (Supplementary video 1C). Besides, considerable variability across the different regions within the same organoid was detected, as the different number of active neurons at multiple neurogenic organoid poles indicated, confirming the impact of regional variability in the neuronal function in brain organoids (Figure 1I). Expression of GCaMP in mature neurons within the organoids was confirmed by immunostaining after recordings (Supplementary Figure 1I).

### Defining the cellular and functional phenotype in Dravet Syndrome brain organoids

As a late-onset developmental disorder, DS organoid modeling has been hampered by the inherent variability in the organoid system. Here, we differentiated brain organoids for 90 days derived from two DS patient-iPSCs carrying the non-synonymous mutation p.A371V:GCT>GTT (DS14 henceforth) and the frameshift mutation p.Val1352CystfrX5 (DS16 henceforth), and a healthy control iPSC line. In addition, we used H9 derived brain organoids as a gold standard differentiation for benchmarking. This maturation stage allowed us to study the DS molecular and functional phenotypes governing the DS epileptogenic state. First, we determined that SCN1A gene expression in control-derived organoids increased significantly in mature stages (day 90 vs day 21 and day 35) (Supplementary Figure 2E). In neurons derived from DS patient iPSC Nav1.1, the sodium channel receptor subunit coded by SCN1A, is preferentially expressed in GABAergic interneurons, but it is also found in a minority of excitatory pyramidal neurons ^27,28^. Unguided brain organoid differentiation generates both excitatory and inhibitory cortical neurons alongside astroglial cells (Figure 2A).

**Figure 2.**
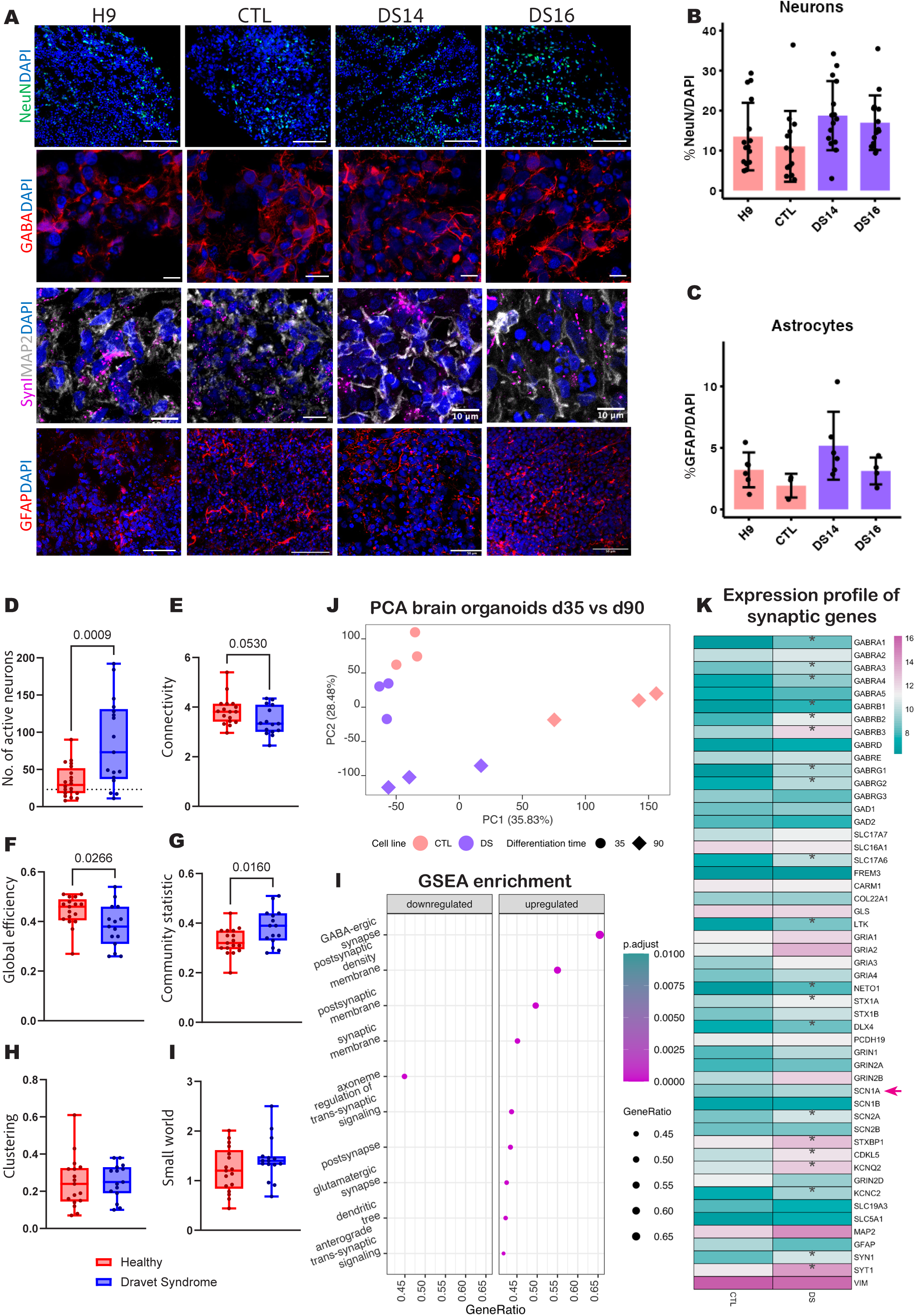
DS brain organoids show imbalanced excitatory-inhibitory circuitry. A. Immunophenotypic characterization of H9, CTL, DS14 and DS16 brain organoids at day 90 of differentiation, showing NeuN, GFAP, GABA, MAP2 and Syn1 markers. B. Percentage of mature neurons (NeuN^+^) in 90-day-old brain organoids from H9, CTL, DS14 and DS16 cell lines. ANOVA test showed no statistically significant differences among cell lines (p value > 0.05). C. Percentage of astrocytes (GFAP^+^) in 90-day-old brain organoids from H9, CTL, DS14 and DS16 cell lines. ANOVA test showed no statistically significant differences among cell lines (p value > 0.05). D,E,F,G,H,I. Calcium signalling analysis of H9 and DS16 brain organoids using GCaMP6s Syn1^+^ neurons. Parameters showing number of active neurons (D), connectivity (E), community statistic (G), global efficiency (G), clustering (H) and small world (I) were calculated per recording. T test was used to test the significance of the results. Only significant p values (< 0.05) are included. J. RNAseq analysis showing late onset phenotype at principal component analysis (PCA) of DS14 and CTL brain organoids at day 35 and 90 of differentiation, distance increased between genotypes at matured (day 90) compared to earlier (day 35) stages. K. Heatmap showing log normalized counts of neuronal and synaptic genes at day 90 of DS14 and CTL brain organoids. Asterisks (*) show differentially expressed (DE) genes between genotypes with p value < 0.05 using Wald test from DESeq2 package. L. Gene set enrichment analysis (GSEA) showing downregulated (left) and upregulated (right) Gene Ontology (GO) categories in DS14 compared to CTL brain organoids both at day 90 of differentiation. No correction was applied for calculating p-adjusted value.

To define the DS phenotype in unguided brain organoids of all conditions were analyzed at 35 and 90 days of differentiation, when neurogenesis is well established, and mature neurons and astrocytes are robustly widespread throughout organoids (Figure 2A and Supplementary Figure 3C). No differences in the pace of differentiation were detected up to day 35 amongst the DS and control genotypes (Supplementary Figure 3A-D). No differences in the proportion of neural progenitor rosettes (Nestin+, SOX2+ and PAX6+) or early born neurons (β3 Tubulin (Tuj1+) were detected by immunofluorescence on day 35 brain organoids. However, transcriptomic analysis suggests that DS organoids mature at a faster pace. The late onset molecular phenotype is evidenced at the transcriptomic level comparing day 35 and day 90 in control and DS (Figure 2 J-F and Supplementary Figure 3 F-G). Yet, day 35 DS organoids showed increased mature neuronal markers (MAP2, SYN1, DLG4, TBR1) gene expression, suggesting an earlier onset of neurogenesis (Supplementary Figure 3 I-J).

Immunophenotypic analysis on the neuronal marker NeuN in DS and control organoids at 90 days of differentiation revealed comparable expression in both groups. Additionally, no noteworthy distinctions were observed in the dendrite marker MAP2, the inhibitory neurotransmitter marker GABA, and the astrocyte marker GFAP across all cell lines (Figure 2 A-C). Yet, these markers showed in all conditions a patched expression throughout the organoid. This intra-organoid variability was confirmed in H/E and NeuN staining in paraffined organoid serial sections (Supplementary Figure 4 A,B), highlighting the intrinsic structural variability in each organoid and confirming the significant intra- and inter-organoid variability independently of the DS and control iPSCs origin.

Next, we evaluated the functional behavior on DS vs control neurons monitoring intracellular calcium levels on 90 days old organoids (DS n=6 and control n=6) expressing GCaMP6s. First, analysis of the calcium signalling recordings showed that DS organoids presented higher number of firing neurons as compared to control ones (Figure 2D). Subsequently, we compared network organization in organoids belonging to each genotype class, as an approximation to a potential epileptogenic *in vitro* phenotype. We used raster plots to compute functional connectivity in groups of the same amount of neurons from both classes (Supplementary Figure 4G). The comparison showed a non-significant trend of reduced connectivity in DS organoids as compared to control ones (Figure 2E). Further analysis of this difference using Global Efficiency, a descriptor that captures the capacity for network-wide communication, showed a better performance in control organoids as compared to DS ones (Figure 2F). This could be related with more segregated networks formed by the neurons in DS organoids, which confirmed by higher community statistic found in this type of organoids, as compared to control ones (Figure 2G). Analysis of clustering and Smallworld parameter in both types of organoids showed no significant differences (Figure 2 H,I).

In summary, DS brain organoids show a circuitry-impaired phenotype, characterized by a reduced whole-network communications, reproducing a late-onset epilepsy. However, this phenotype is not detectable at a cellular subtype and architectural level.

### Transcriptomic analysis indicates an excitatory-inhibitory circuit imbalance in DS brain organoids

DS, as many other DEEs, show the first clinical symptoms postnatally, during the first two years of life, when the human cerebral cortex is building its circuitry ^29^. Thus, we further delved into the transcriptomic phenotype of maturing organoids at day 90, when synaptogenesis is increasing (3 samples from 2 independent differentiations).

Upon exploring the sample distribution using dimensionality reduction (PCA), noticeable distinctions were observed, particularly at mature time-points. During these stages, DS and control brain organoids were distinctly separated along the principal components (Figure 2J). At day 90, 1100 genes were differentially expressed between control and DS organoids and at day 35, 360. Sixty-four of them (17.8%) appeared as DE in both 35 and 90 days (Figure Supplementary 3E). Interestingly, these 64 genes followed the same expression trend at both timepoints. DE genes between early and mature stages, including those common to both stages, were involved in basic neurodevelopmental functions, such as cell junction assembly and synapse formation (Figure Supplementary 3H). Together, these data suggest that the molecular phenotype related to regulation of synapse structure or activity is triggered already at early maturation stages (Figure Supplementary 3G).

Delving deeper into the molecular phenotype at day 90, 663 out of 1100 genes were upregulated and 447 were downregulated between the control and DS organoids. Gene Set Enrichment Analysis (GSEA), ranked according to their significance (p-adjusted value), indicated that the most significantly dysregulated biological processes were associated with essential neuronal homeostatic functions linked to synapse formation and organization (Figure 2J-L). Among genes differentially upregulated in DS organoids, a remarkable correlation was found with genes driving other epileptic syndromes, such as SYN1, STX1A, STXBP1, KCNC2, all involved on neuronal excitability, connectivity and brain plasticity mechanisms. Instead, SCN1A, DS pathogenesis driving gene, did not show differential expression (Figure 2K). Moreover, DS showed a generalized trend to reinforce the excitatory synapse activity (Figure 2K). Also, there is an upregulation of GABA receptors (Figure 2K), expressed in excitatory neurons ^30^, suggesting a putative compensatory mechanism to the limited inhibitory synaptic activity in DS.

No significant changes in cortical excitatory neuronal marker, such as TBR1, FOXP2, SATB2, EMX2 and EMX1 neither significant downregulation of astrocytic markers, such as GFAP and vimentin, was detected (Figure 2K), supporting the immunophenotypic data. These results indicate that there is an affectation in both the excitatory and inhibitory synaptic signalling and connectivity that disrupts its normal electrophysiological equilibrium.

### CNN integrated brain organoids phenotype classification

In DS patients and organoids, the pathogenic phenotype relies on an abnornal axonal circuitry. Thus, the determination of the phenotype over the intrinsic variability results complicated with conventional immunofluorescence markers. To overcome the limitation of evaluating in real-time changes in the circuitry in a genotype related backgrounds, we leveraged Cellbrite staining and live imaging.

Thus, we aimed to determine whether the DS-associated phenotype, involving the neuronal axonal network, can be systematized to overcome the limitations due to the intraorganoid variability and late onset phenotype.

To this purpose, we leveraged the axonal tracer capacity of Cellbrite staining to label the axonal network *in vitro* and establish an integrated protocol for live image acquisition of the whole organoid rather than histological sectioning. To efficiently boost the potential of the live image analysis in Cellbrite stained organoid, we integrated a multi-step deep learning (DL) pipeline (Figure 3A). Whole Cellbrite stained organoids were imaged in depth (∼500 µm) using wide-field microscopy at low magnification. This step was followed by extensive pre-processing, encompassing the following serial steps: 1) whole surface stitching considering all z-stacks, 2) intensity and pixel (px) sharpness correction, 3) partition of stitched corrected image into 224x224 px, and 4) removal of non-QC images based on px focus sharpness, artifacts due to reconstruction process and amount of background, resulting in corrected organoid region images (CORIs). Henceforth, the integrated pipeline and analysis will be named Image Phenotype Network (ImPheNet). All CORIs were analyzed independently, namely, the ImPheNet system outputs a prediction per each CORI (ranging from 0 DS to 1 for control). The performance of the model reached an accuracy of 0.732 (Supplementary Figure 4C). As emphasized earlier, the variability within organoids is substantial. Therefore, we consolidated the CORIs prediction value into an adjusted classification score for each whole organoid estimated from the training dataset (See Methods Section for details). These scores are named as CONTROL SCORE and DRAVET SCORE, resulting in 0 = totally Dravet and 1 = totally Control (Figure 3 B-D). Using the classification score, the phenotype identified matched with the genotype with an accuracy of 0.967 (Supplementary Figure 4D).

**Figure 3.**
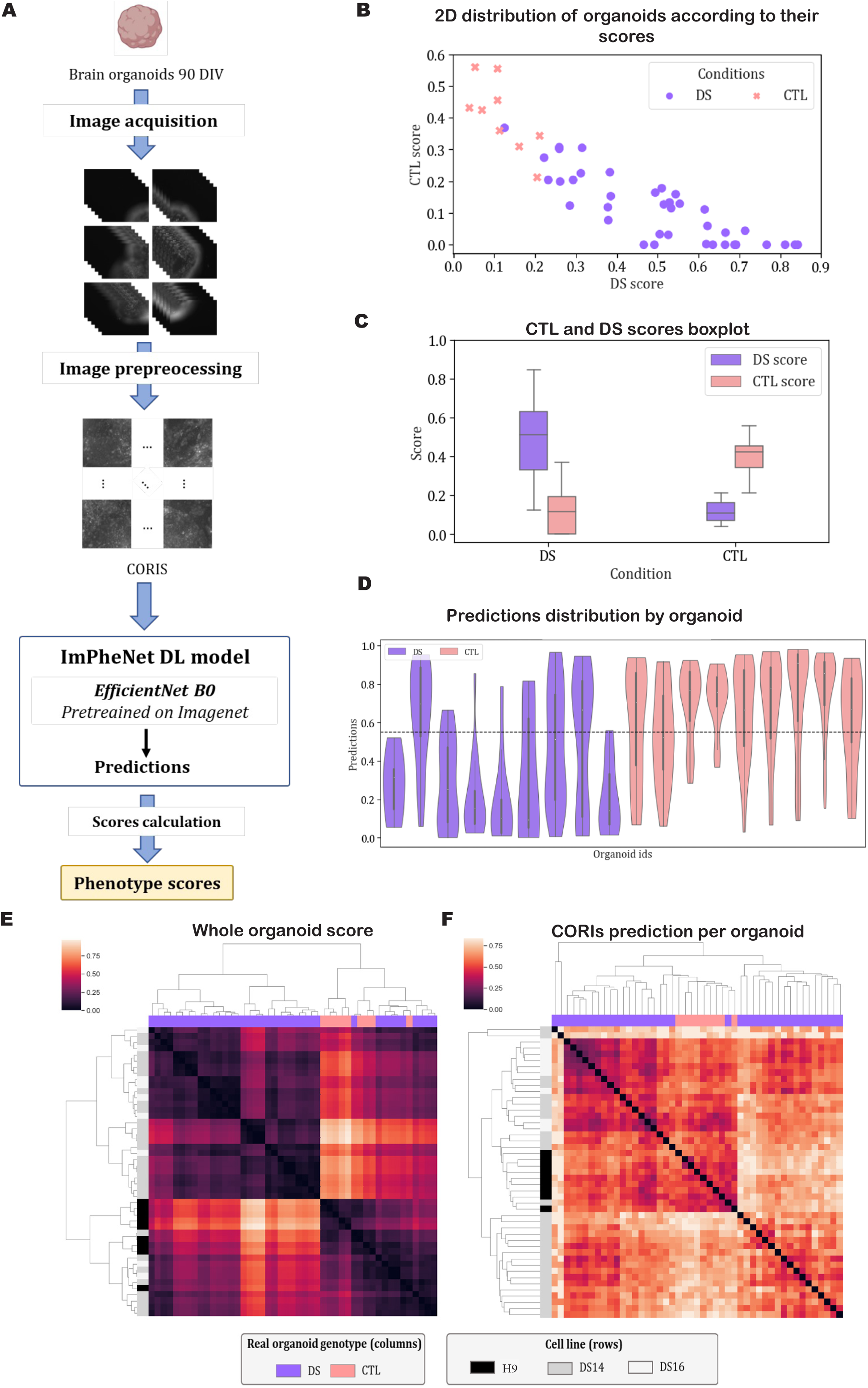
ImPheNet predictive model. A. Workflow of ImPheNet AI-based model: from image preprocessing to genotype-related phenotype prediction. B. Representation of “CTL score” (y axis) and “DS score” (x axis) per organoid, labeled based on their real genotype (H9 vs DS14). C. Boxplot of CTL and DS score ratios per genotype. D. Violin plot of phenotypic predictions (CORIs) per organoid. E,F. Clustermap using the Jensen Shannon distance (left) between integrated whole-organoid histograms of predictions and (right) clustermap using the Euclidean distance between the computed scores, per each CORI grouped per organoid. These heatmaps show how the predictive power dramatically increases when whole organoid is considered regarding to consider only regional information, highlighting the need of an integrated organoid analysis, to reduce the intra-organoid variability bias.

The intra-organoid variability is a confounding factor when it comes to predict the organoids phenotype. This variability is reflected in the dispersion observed in the CORIs prediction values in the same organoid (Figure 3D and Supplementary Figure 4D). When comparing the data distribution of CORIs classification and whole organoid adjusted scores, the dissimilarity among each organoid belonging to the same class is reduced (Figure 3E left). Notably the variability due to batch effect is buffered using the ImPheNet algorithm independently of whether the classification is carried out on whole scores or on single image classification (CORIs) (Figure 3E right). Altogether, this analysis indicates a robust DS phenotype detectable at live imaging levels enhanced by ImPheNet classification, correlating with the phenotype detected at a transcriptional level.

### ImPhenet classification of Anti-seizure drugs toxicity on DS brain organoids

In DS, as in many other rare disorders, limited randomized controlled trials have hindered evidence-based treatments, the only being available are for the use of stiripentol (STP), cannabidiol (CBD), and fenfluramine (FFA). Treatment guidelines recommend valproic acid (VPA) as the preferred first-line treatment ^31^. Sodium channel blockers, such as Phenytoin (PHT) and Lamotrigine (LAM) are not recommended due to negative effects on seizure frequency and cognitive outcomes ^32^. VPA can reduce seizure frequency and severity, though achieving seizure freedom is rare. Second-line options outlined in the guidelines include STP with VPA and clobazam, topiramate, and the ketogenic diet. Due to the potential negative effects of antiseizure drugs in DS patients, we aimed to explore the impact of other known antiepileptic treatments in ImPheNet as a tool for drug testing in DS brain organoids. All treatment conditions were applied acutely for one week in mature brain organoids (day 90). Cellbrite stained organoids were imaged pre- and post-treatment and no evident differences in the morphology nor in fluorescence intensity amongst them were detected upon treatment (Figure 4A). Subsequently, the ImPhenet pipeline was used as previously described. Two cohorts of brain organoids were generated for the training dataset: DS untreated brain organoids and those treated with the sodium channel blockers Lamotrigine and Phenytoin (Na INH class), serving as markers of toxicity-related outcomes. Once the model was trained and validated, we further tested the potential toxicity of VPA, CBD and FFA in DS brain organoids.

**Figure 4.**
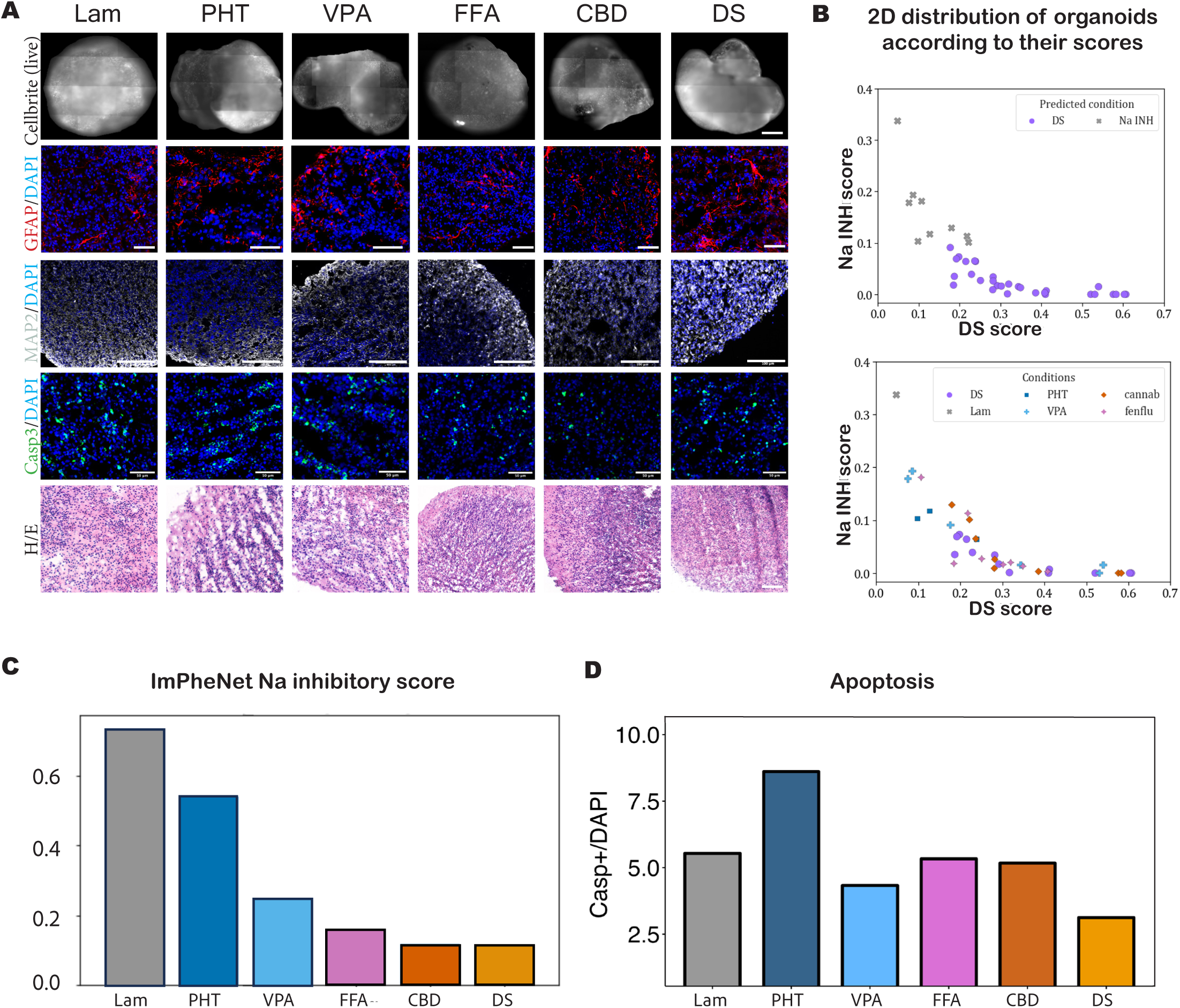
ImPheNet prediction in anti-seizure treatments. A. Morphological and immunophenotypic analysis of antiseizure drug effect on DS day 90 brain organoids Cellbrite stained organoids imaged after 7 days of treatment (Row 1). Immunophenotypic (astrocytes-GFAP; neuronal dendrites-MAP2; apoptosis-activated CASPASE3) characterization (Rows 2-4) and Hematoxylin and eosin (H/E) staining (Row 5) of brain organoids exposed to Lamotrigine (Lam), Phenytoin (PHT), Valproat (VPA), Fenfluramine (FFA), Cannabidiol (CBD). These results indicated limited morphological and cellular changes. B. 2D distribution of DS and Na INH scores per organoid colored by predicted class (up) and treatment (down). C. Histogram showing Na INH score of the different treatments in DS brain organoids calculated with ImPheNet. Na INH score is represented by the ratio of the NA INH CORIs. D. Percentage of activated Caspase-3 cells in drug-induced apoptosis analysis by IF staining. ANOVA test revealed no significant differences among treatments (2 independent experiments/ n=3 organoids per condition (p value > 0.05).

During algorithm development, it was detected that adjusted classification scores resolved better the classification on toxic treatment events than CORI’s predictions (CORI’s-Accuracy 0.759 vs whole organoid score accuracy 0.941) (Figure 4B,C and Supplementary Figure 5B,C). Notably, recently approved drugs for DS, CBD and FFA, exhibited reduced levels of the Na INH class component, followed by VPA (Figure 4C), suggesting that they show lower levels of toxicity. Transcriptomic analysis on VPA treated DS and control organoids reveals that the anti-seizure drug induced significant changes in both genotypes (Supplementary Figure 5F,G). These results were validated in parallel using organoid histological analysis, encompassing neuronal and glial markers, alongside activated caspase 3 (aCasp3) immunostaining to determine putative increase in apoptosis (Figure 4A, D). No significant changes could be detected neither on the cell type presence nor apoptosis in the treatments, suggesting that Na INH class drugs can affect functionally to the neuronal network rather than to the cytoarchitecture in the brain organoids. Altogether, the ImPheNet rises as a relevant tool to detect toxicity events in DS-derived brain organoids, that cannot be detected by conventional immunostaining methods.

## DISCUSSION

In this study we propose a systematic DL-driven analysis, the ImPheNet, to overcome the brain organoid variability generated with unguided differentiation protocols. Amongst the advantages of the use of brain organoids is the intrinsic generation of a neuronal network that allows to model disorders affecting the connectivity. The ImPheNet allowed us to perform live imaging classification of DS-derived organoids, the developmental epileptic encephalopathy we employed as a paradigm of the late onset pharmacoresistant neurodevelopmental disorder. Moreover, we benchmarked the DL detected phenotype with molecular and functional analysis that determined the unbalanced inhibitory-excitatory circuitry nature of DS.

### Intra- and inter-organoid variability

The use of brain organoids in disease modeling and translational studies has been hampered by the substantial variability inherent to the model. Brain organoids differentiation protocols start with a defined number of self-aggregating iPSC or hESC, to enable correct differentiation. Upon neural induction, multiple neurogenic centers or rosettes are generated ^33^. In unguided organoid differentiation protocols, the wide range of the brain cellular diversity is generated following clonallity of the NPC clusters distributed throughout the organoid neuroepithelium at very early stages ^11^. Thus, this regional patterning segregation occurs as early as day 15 of differentation, making organoids regional variablity very liable to small signalling variations in the culture media, and other deviations in the protocol (stemness status, survival to seeding, small molecule bioavailability, and morphological arrangement) ^33^. The ability of brain organoids to grow for several months in suspension, enables long-term reconstruction of a complex neuronal network that rearranges with time ^14^. Moreover, the suspension nature of the 3D model growth impairs the systematic positioning of the organoids for longitudinal analysis, increasing the morphological variability perception. In this study, we tackled the contribution of the neuronal network and its functional capacity to the given inter- and intra-organoid underlaying brain organoid variability. Staining axonal networks with Cellbrite, a DiO derivative, and Venus reporter lentiviral low dosage infection, we determined that brain organoids axons build bundles to functionally connect far-distance regions, risen from multiple neurogenic centres, similarly to other protocols using air-liquid interface ^14^. The non-homogenous distribution of the neuronal connections and their constant remodeling as the organoid matures constitute a great source of the observed organoid variability. Using calcium imaging, we did not observe genotype-related differences in the transmission of the information across different regions considered separately, although we can distinguish reduced global efficiency in DS compared to control organoids when considering all regions grouped per organoid. This strong reduction in inter-organoid variability makes organoids from the same genotype comparable between them and distinguishable from the others in the same way the ImPhenet does, and thus, highlights brain organoids as excellent tool for brain network analysis and disease modelling.

### Integrative Method for Brain Organoid Phenotyping

AI methods, including Convolutional Neural Networks (CNN), are especially suited to analyze complex data derived from brain organoids ^34–36^. Live imaging of medium-large 3D densely populated structures encompass a good number of optical challenges, including size, systematic position, light refraction and penetrance. Here, we propose a reductionist approach using Cellbrite dye to label neural membrane followed by live imaging on a wide-field microscope. Using the ImPhenet, we determined that only when brain organoids are contemplated as a whole, instead of as the mean of multiple regions, DS phenotype arises. The ImPheNet scoring algorithm significantly reduced the inter-batch variability, providing a robust analysis methodology for further applications, such as *in vitro* drug testing.

DS patients are pharmacoresistant. Consequently, there is a compelling need to enhance their treatment alternatives. Unfortunately, to date, only a few randomized controlled trials of new drugs have been performed ^31,37^. Not even common antiseizures sodium channel antagonists drugs, such as lamotrigine or phenytoin, have been systematically analysed in DS, resulting in uncontrolled seizures ^38^. In this study, we provide a proof-of-concept that the ImPhenet algorithm can be leveraged to evaluate *in vitro* the toxic potential of antiseizure drugs. The ImPhenet algorithm classifies the recently approved drugs, CBD and FFA, as the less toxic in DS, followed by VPA, the first line and most commonly used drug in DS treatment and other epileptic encephalopathies. Although more data is needed in this context, with further drugs and more patient derived iPSC, this is a promising result to advance in the therapeutic control of epileptic encephalothies.

### Brain organoid modeling of DS

The primary genetic cause of DS is the presence of mutations in the SCN1A gene, which encodes a subunit of the sodium channel Nav1.1 ^39^, expressed in inhibitory neurons. SCN1A mutant mouse models have shed light on the cell types implicated in DS ^40–42^. However, not a full neuropathology reconstruction has been achieved, mainly due to inherent evolutionary differences in the human brain construction, relating to the multiple types and amount of neurons as well as to the higher complexity of the human neural network and synapse plasticity ^43–47^. Our data obtained from DS brain organoids indicate, that at a transcriptomic level, the DS organoid phenotype is strongly associated to synaptic homeostasis and function, and not to the cellular subtypes. Notably, these differences were translated into changes at the circuitry level, where we identified significant changes in the construction of the neuronal network, pointing towards and imbalance in the excitatory-inhibitory network. Although at 90 days of differentiation, collective events were not detectable by intracellular calcium recordings, differences between control and DS organoids were notable in the developing networks. Higher numbers of active neurons in DS organoids were not accompanied by increased global efficiency. In contrast, higher segregation observed in DS organoids as compared to healthy organoids resulted in less efficient networks, which may lead to increased excitation at later timepoints. Our data add to that from multiple labs specialized in neurodevelopmental disorders iPSC modelling, supporting the relevance of the alterations in the neuronal circuitry ^4,6,22,48^. Altogether, our data support expand on the use of brain organoid modeling to complex neurodevelopmental disorders involving circuitry unbalance, and overcomes the limitation of the organoid use by establishing a whole-organoids system for the genotype-related phenotype analysis.

### CONCLUSIONS

We have developed a novel approach to analyze phenotypes related to neuronal axonal network changes using a reductionist and automated way, the ImPheNet, that it is compatible with longitudinal brain organoid studies, like for instance *in vitro* drug testing. Moreover, we provide comprehensive study of the source of the variability in brain organoids and we propose a tool to overcome it. Last, we provide insight of the neuropathogenesis of DS and the molecular effect of VPA as a DS antiseizure drug.

## STAR METHODS

### Lead contact

Further information and requests for resources and reagents should be directed to and will be fulfilled by the lead contact, Sandra Acosta

### Materials availability

This study did not generate new unique reagents.

### Data and code availability

RNA-seq data have been deposited at GEO and are publicly available as of the date of publication. Accession numbers are listed in the key resource table GSE256142.

All original code has been deposited at Github and it is publicly available as of the date of publication.

Any additional information required to reanalyze the data reported in this paper is available from the lead contact upon request.

### Experimental Models and Methods details

#### Brain organoids generation

Commercial embryonic stem cell line WA009 (H9), iPSC control (CW70019), and commercial iPSCs derived from Dravet Syndrome patients (Ref.PFIZi014-A, PFIZi016-A, EBISC) were cultured in mTESR1 (Ref.85850, Stem Cells Technologies) on 1:40 Matrigel (Ref. 45354277, Cultek) until they reached 70-80% confluence in 6-well plate multiwell format. To generate the brain organoids following a previously published protocol {Lancaster_2014}, cells were dissociated using Gentle Cell Dissociation Reagent (Ref. 100-0485, Stem Cells Technologies)and plated in mTESR1 with Y27632 Rock Inhibitor (TO-1254-10 MG, Biogene) in 96 round bottom non-adherent cell culture plates (Ref. CLS7007-24EA, Corning) at a density of 9000 cells per well. Once self-aggregated organoids reached a diameter of 500-600 µm, approximately on day four of differentiation, they were switched to neural induction medium, as in Lancaster, 2014^8^ .

Upon the formation of the neuroepithelium, which is visible at approximately day 8-10 depending on the iPSC line, the organoids were embedded in a Matrigel droplet and transferred to non-treated 6mm plates with cerebral organoid differentiation medium without vitamin A. The neuroepithelial buds were allowed to expand in this medium for four days under static culture conditions. After four days, the embedded organoids were transferred to orbital agitation with Vitamin A supplemented cerebral organoid differentiation medium for further maturation, as described in Lancaster et al. 2014. Brain organoids were matured on agitation (70rpm) for 90 days at 37°C and 95% humidity, with partial media replacement twice a week.

#### Drug testing experiment

Ninety-days old organoids were accutely treated for one week with antiseizure drugs, including Valproic Acid, Phenytoin, Lamotrigine, Cannabidiol, and Fenfluramine. Briefly, three organoids per well were plated into a 24-well plates (Ref.142485, DDBiolab), and treated with the anti-seizures drugs (final concentration VPA-50mM (HY-10585, Quimigen), PHT-1mM, Lam-200µM, CBD-10µM, and Fenfluramine-10µM). Half-volume medium change,maintaining drug concentration, was performed on day three. For live- imaging, organoids were stained with Cellbrite (0.5uL in 100uL of media) dye for 1 hour under agitation, and imaging was performed before starting the treatment and after 7 days, prior to fixation.

#### Immunofluorescence Analysis

Cerebral organoids were fixed for 1 hour at room temperature with 4\% paraformaldehyde (PFA) and washed three times with 1x PBS (phosphate-buffered saline) before overnight cryopreservation in 30% sucrose. After cryopreservation, organoids were embedded in OCT (05-9801, Bio-Optika), and cryosectioned at 10 µm-thick slices using a Leica CM1860 UV cryostat. For immunostaining, sections were Blocked/permeabilized in buffer solution containing 3% donkey serum (D9663, Merck), 2% bovine serum albumin (BSA)(A2153, Merck) and 0.1% Triton X-100 (X100, Merck) in 1x PBS for 1 hour at room temperature. Subsequently, sections were incubated overnight with primary antibodies at 4°C (See Supplementary Table 1 for details)Table\ref{Table1}. Sections were washed twice 1x PBS containing 0.1% Triton X-100 (PBS-T), stained with secondary antibodies, washed 3x in PBS-T and labeled with DAPI for nuclear counterstaining. Slides were mounted in Mowiol® 4-88 (31381, Merck). Imaging was performed using Zeiss LSM 880, Zeiss LSM 980 Airyscan2 inverted confocal microscopes, or Zeiss Axio Imager M2 Apotome 3. For image processing and quantification, FIJI (Image J) was used. Brightness was adjusted for each maximal projection individually according to z-stack volume. Quantifications were performed using Fiji’s Analyze functions: particles tool involved manual intensity correction before binary conversion and Watershed segmentation. Size and circularity parameters were adjusted depending on the image magnification and the nature of the staining (nuclear or cytoplasmic). 3D viewer (Imaris) was used to record the video shown in Supplementary video 1A.

#### RNA preparation

Brain organoids were harvested and snap-frozen in liquid nitrogen and stored at -80°C upon their processing. RNA extraction was performed as followed: mechanic homogeneization in 700 µl Qiazol (Qiagen), phase separation in 140 µl Chloroform (C2432, Merck) followed by 15 minutes centrigution at 12000rpm; precipitation in Isopropanol (19516, Merck) and rehydration in 70%, final dilution in 100 µl of ultraPure water. Next, a column purification with the RNeasy Miki Kit (Qiagen) was performed following the supplier protocol. QC was validated in a Bioanalyzer.

For each condition, the following experimental approach was determined: CTL day 21 (2 replicates; 6 organoids per replicate (OxR)), day 35 (2 replicates, 4 OxR), and day 90 (2 replicates, 2 OxR); iPSC DS and control day 35 and day 90 (3 replicates, 2 OxR); and day 90 organoids after VPA treatment from both control and DS lines (2 replicates, 2 OxR).

#### RNAseq Analysis

RNA sequencing was conducted at the Universitat Pompeu Fabra (UPF) Genomics Service. RNAseq libraries were prepared using polyA capture and QC validated in Bioanalyzer. Libraries were pooled together, and sequenced on a NexSeq 500, using a 2x75 cycles High Output run, in addition to a 2x75 cycles Mid Output run. Transcript alignment was performed on STAR software was utilized against the human reference GRCh38 and annotation version 109 from Ensembl. Counts were obtained using HTSeq software, and MultiQC software was employed for quality assessment. The count matrices were fed into the DESeq2 statistical package (R) to determine differential transcript expression between samples, employing adjusted p-value and lfcThreshold parameters of 0.05 and 0.58, respectively. DESeq2 was also used to generate a principal component analysis (PCA) plot, illustrating the variance between distinct sample groups and the similarity within sample replicates. Distance heatmap was generated to display the large distance between time points and the tight clustering within the same replicates. For the visualization of heatmaps and other visual representations derived from the analysis of differential expression, ggplot was implemented. To conduct the Gene Set Enrichment Analysis of Gene Ontology, the gseGO function from clusterProfiler (Wu et al. 2021) was employed.

#### H&E and IHC

For hematoxylin/eosin staining, fixed and cryosectioned samples were subjected to hematoxylin staining, which was carried out for 5-10 minutes. To enhance nuclear contrast, the slides were submerged in water for 5 minutes before incubating with eosin staining for 1 minute to highlight cytoplasmic structures.Finally, the samples were mounted for microscopic examination.For inmunohistochemistry, EnVision Flex kit (GV80011-2, Agilent) was used through DAKO Autostainer. Briefly, fixed and cryosectioned samples were incubated in antigen retrieval solution (EnV FLEX TRS, High pH) for 30 minutes at 97°C prior to 20 minutes incubation with primary antibody (NeuN policlonal, Novusbio). After, the sections were washed, incubated with the blocking solution (EnV FLEX PeroxidaseBlocking Reagent) for 3 minutes and finally incubated with an HRP conjugated secondary antibody (EnV FLEX/HRP) for 20 minutes.

#### Calcium signalling analysis

Live calcium imaging was performed in 6 control (H9 hESC line) and 6 Dravet (DS16 iPSC line) brain organoids at day 90 of differentiation using the GCaMP6s calcium indicator. GCaMP6s was expressed in neurons using AAV7m8-Syn1-GCaMP6s. For the infection, organoids were transferred into a 48WP. For 4 organoids, 2.5 uL of the virus was diluted in 400 uL of differentiation media. The day after, 400ul of differentiation media was added to each well. After that, half of the media was changed twice a week as usual. Intracellular calcium imaging was performed in an Olympus IX70 fluorescence inverted microscope equipped with a fast Hamamatsu camera using 10x objective and 33 frames per second. Several regions of each organoid were recorded during 11 minutes at days 7, 11 and 14 after infection in BrainPhys media. Recorded movies of intracellular calcium fluorescence were analyzed with NETCAL software (www.itsnetcal.com) ^49^ in combination with other custom-made packages, providing the raster plots of spontaneous neuronal activity and connectivity matrices. Neural interactions and functional connectivity were captured using Global Efficiency and Community Statistic as previously described ^50,51^

#### Low dosage viral infection

Second generation lentivirus production was performed by transient transfection of 293T cells with a packaging vector (psPAX2), an envelope vector (pMD2.G) and the lentiviral vector pV2luc2, which contains a Venus reporter cassete under the control of an ubiquitous promoter. After 48 and 72 hours of transfection, supernatants containing viral particles were collected, pooled, and rapidly frozen at -80°C for storage.

90 days-old brain organoids were transduced using 1:5 and 1:20 diluted supernatant in maturation media. After a 24-hour infection period, brain organoids were washed and supplied with fresh maturation media. 3D viewer (Imaris) was used to record the video shown in Supplementary video 1B. SNT pluggin from FIJI was used for the segmentation of images shown in Supplementary Figure 4.

#### Live imaging

For live imaging, matured brain organoids at day 35 and day 90 were incubated with green Cellbrite (0.5uL in 100uL of media) dye for 1 hour under agitation. Throughout the imaging process, the organoids were kept in BrainPhys Imaging Optimized Medium. Wide-field imaging was performed using a Zeiss Axio Observer Z1 microscope with a 10x objective and stitching option (10% overlap). Images were captured in a z-stack with 50µm intervals. The stitched images were then exported in .czi format, and the separated tiles were saved in .tiff format for subsequent analysis. For viral transduced neuronal tracing, organoids were transferred to a 24-well iBIDI plate with BrainPhys media and subjected to imaging using a Confocal Carl Zeiss LSM 880 at 37°C with CO2 settings. Images were captured at 25x magnification with oil immersion.

#### Image Preprocessing

All the images from the dataset have been preprocessed before using them to train the DCNN following the steps bellow described. Focus stacking(1): to generate a suitable 2D projection for each stack of images representing the same x and y pixels but at different depth we have implemented an algorithm based on the Laplacian of Gaussian (LoG). This algorithm has been implemented using the Laplace and gaussian filter offered by the skimage package in Python using a σ=2 and a kernel size of 10 for the gaussian filter and a kernel size of 10 for the Laplacian filter. Since the regions on focus are regions of rapid intensity change, the pixels in these regions will get large LoG scores. Nevertheless, the regions out-of-focus are blurry and the pixels in these regions will get low LoG scores. Thereforeonly the intensity value at the stack where the LoG (x, y, z) achieves maximum is retained. Image intensity correction (2): minimum and maximum intensity values from all the tiles representing the same organoid are considered to homogenize the intensity of the different images by applying a first normalization step. Then, pixels with an intensity value over 25 are selected to define the organoid region. Those below the threshold are considered as background. Intensity gamma correction according to the mean intensity value of the organoid region is applied to the image in order to homogenize the intensity among tiles (the gamma values used for the correction are 0.6 if the mean intensity value is bellow 25, 0.8 if the mean intensity value was between 25 and 65, 1 if the mean intensity value was over 65). Image stitching (3): ImageJ plugin Grid/Collection Stitching is used to stitch the different tiles and generate a single 2D image. Phase correlation and fast Fourier Transform are applied to find the correct overlap of connected tiles, and linear blending fusion algorithm is used to ensemble the different tilesin a single image. Image fragmentation (4): A window filter of 224x224 with a stride of 224 was used to fragment each pre-processed image. Fragments with at least 90% of pixels over the intensity threshold (> 35) were selected and included in the dataset. Data clearing (5): Finally, manual filtering is performed to remove non-QC images Filtering paramenters include: images of the external part of the organoid containing allos, artifacts caused by image reconstruction process, blurred images and background images not discarded in the previous step.

#### ImPheNet: Dataset generation

For phenotype prediction, it is generated a dataset containing CTL (iPSC CTL and hESC H9) and DS (DS14 and DS16) images acquired from 90 days-old brain organoids after 1h of Cellbrite incubation (from a total of 100 organoids). A second dataset for drug testing organoids was created with the following classes: treated (INH) and untreated DS (DS14 and DS16) 7 days after Cellbrite incubation. CORIs derived from the same organoid were labeled with the same ID and allocated to the same subset of data (training-validation-test). In both cases, 75% of CORIs were used for training the model (incluing 25% for model validation) and 25% for testing. Stratified splitting based on organoid ID ensured that each subset kept the same proportion of classes than in the original dataset. However, since not all the organoids contain the same amount of CORIs the proportionality between classes is just partially respected.. Data augmentation based on image rotations and image flips was implemented in an online approach to enlarge the dataset. Intensity distribution in each CORIs was corrected using the gamma correction factor (based on mean intensity). Finally, a normalisation step based on histogram stretching was applied to ensure that all the images are defined by the same range of intensity values (from 0 to 1).

#### ImPheNet: DL model

Two different DL models were trained considering the question to address: one for genotype prediction (identification of organoid genotype through its phenotype) (1) and another for drug testing (Identification of organoids treated with sodium channel inhibitory drugs) (2). Respectively, the output value of each model are: the probability of a particular region of being control (1) and the probability of a particular region of being affected by a sodium channel inhibitory drug (2). Both models were generated by transfer learning from pretrained models available in Python Tensorflow library and using different combinations of hyperparameters. Each final model was selected according to validation accuracy (the highest the best) and validation loss (the lowest the best). The specifications for each selected model are described in Table 1.

**Table 1.**

| Antibody | Company (reference) | Description |
| --- | --- | --- |
| Anti-TUBB3 (Tuj1) mouse | BioLegend (801201) | 1:1000, 4°C O/N |
| Anti-GFAP, chicken | Merck (AB5541) | 1:1000, 4°C O/N |
| Anti-GFAP, rabbit | Genetex (GTX108711) | 1:500, 4°C O/N |
| Anti-MAP2, chicken | Abcam (ab5392) | 1:1000, 4°C O/N |
| Anti-Nestin, rabbit | Biolegend (841901) | 1:500, 4°C O/N |
| Anti-PAX6, rabbit | Genetex (GTX113241) | 1:500, 4°C O/N |
| Anti-SOX2, rabbit | Invitrogen (48-1400) | 1:500, 4°C O/N |
| Anti-Pro-Caspase-3 | Sigma (C8487) | 1:5000, 4°C O/N |
| Anti-NFM, rabbit | Abclonal (A19085) | 1:500, 4°C O/N |
| Anti-GABA, rabbit | Sigma (A2052) | 1:500, 4°C O/N |
| Anti-Syn1, rabbit | Novus Biological (NB300-744) | 1:500, 4°C O/N |
| Anti-NeuN, rabbit | Abcam (ab177487) | 1:500, 4°C O/N |
| Anti-NFH, rabbit | Genetex (GTX110065) | 1:500, 4°C O/N |
| Anti-GFP, mouse | Invitrogen (11559046) | 1:500, 4°C O/N |
| Anti-rabbit, AlexaFluor 488, donkey | Thermo (A-21206) | 1:1000, 4°C O/N |
| Anti-mouse, AlexaFluor 594, goat | Jackson (715-585-150) | 1:500, 4°C O/N |
| Anti-chicken AlexaFluor 647, donkey | Jackson (703-605-155) | 1:1000, 4°C O/N |

For each model, all the metrics (accuracy, jaccard similarity, dice similarity, sensitivity, and specificity) were calculated using the SciPy python package considering each sample independently. For detailed information about the model see Supplementary Table 2.

#### ImPheNet: DL model results

The binary outputs of the DL models were transformed to a score classification using a simple mathematical computation. First, all the CORIs predictions from the same organoid were joined in a vector that was then split in two considering the value of the predictions: predictions below and above a given threshold.The threshold was settled using the validation subset and selecting the value which maximazes the validation accuracy. Secondly, all the predictions were scaled from 0 to 1 to get the class intensity of all the samples in the same scale (Eq 1). This is necessary in case the threshold is not 0.5. Then, the mean intensity per class is obtained and the class score is computed as in Eq 2.

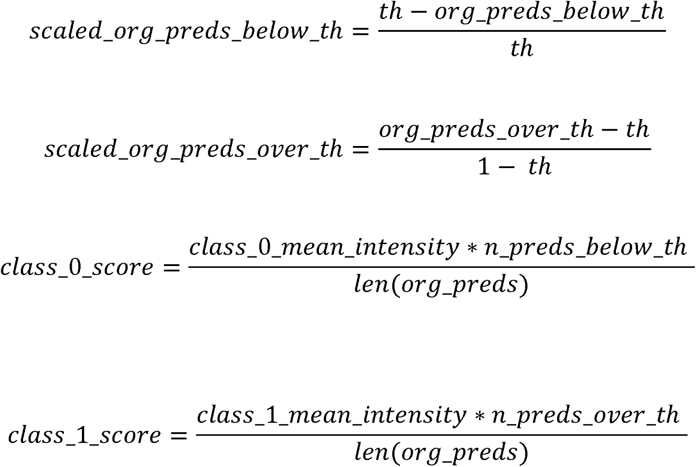

As a result of the scoring, we generated two statistics. In the case of the phenotype DL model, class 0 correspond to DS and class 1 to CTL. In the case of the drug testing DL model, class 0 correspond to DS organoids in control conditions (non-treated) while class 1 correspond to DS organoid treated with a sodium channel inhibitory drug. For the representation of the predictions from all CORIs coming from the same organoid, it is used a histogram, specifically a normalised histogram of 20 bins. Each histogram bar reresents the amount of predictions comprised in each subrange of the probability range a normalized histogram of 20 bins was created counting the number of predictions comprised in each region of the probability range (0-1 divided in 20). Histogram normalisation is key to compare the results coming from organoids with different number of samples.

**Supplementary Table 2.**

|  | **DL model for phenotype prediction** | **DL model for drug testing** |
| --- | --- | --- |
| *Pre-trained architecture* | EfficientNet B0 pretrained on ImageNet | EfficientNet B0 pretrained on ImageNet |
| *Number of unfrozen layers* | 1 fully connected layer with 16 neurons + 1 fully connected with 1 neuron (classification layer) | 1 fully connected layer with 16 neurons + 1 fully connected with 1 neuron (classification layer) |
| *Initial learning rate* | $2.5 \cdot 10^{-3}$ | $5 \cdot 10^{-4}$ |
| *Input image size* | 224x224x3 | 224x224x3 |
| *Batch size* | 93 | 71 |
| *Class weights* | None | DS weight: 0.74 / INH weight: 1.53 |
| *Optimizer* | Adam with default parameters | Adam with default parameters |
| *Regularization techniques* | Dropout of 0.4. Batch Normalization, Early stopping (stop if not validation improving after 5 epochs), Learning rate decay: $\text{new\_lr} = \text{current\_lr} - 0.5 \cdot \text{current\_lr}$ each 2 epochs without validation loss improvement | Dropout of 0.4. Batch Normalization, Early stopping (stop if not validation improving after 5 epochs), Learning rate decay: $\text{new\_lr} = \text{current\_lr} - 0.5 \cdot \text{current\_lr}$ each 2 epochs without validation loss improvement |
| *Activation functions* | Last layer: Sigmoid which offers a “probability” of belonging to a particular class. / Other layers: Rectified Linear function (ReLU) | Last layer: Sigmoid which offers a “probability” of belonging to a particular class. / Other layers: Rectified Linear function (ReLU). |
| *Loss function* | Binary cross-entropy | Binary cross-entropy |
| *Objective function* | Accuracy | Accuracy |

## ACKNOWLEDGEMENTS

We thank Dr. Oliver, Dr. Bertranpetit, Dr. Fiore, Dr. Alcántara and Dr. Ortega for their insights in manuscript. We thank IDIBELL and UB Scientific Platforms, especially to Dr. Torrejón at Bellvitge-Microcoscopy Unit CCTIUB, for their constant technical assistance in the experimental execution and the UPF Genomics Platform for the execution of RNAseq. S.A. is a Serra-Hunter Assistant Professor Fellow of the Generalitat de Catalunya. This project has been funded with Caixa Impulse (CXI20-0002) to S.A. and O.L, AGAUR Llavor (2021) to S.A., Ministerio de Ciencia y Innovación (MICINN) (PID2021-128208NB-100) to S.A. This work has received funding from the European Union’s Horizon 2020 research and innovation programme under the grant agreement NeuChiP No. 964877 (J.S. and D.T.). E.E is funded by a Ramon y Cajal Fellowship (MICINN).

## Notes

### Competing Interest Statement

The authors have declared no competing interest.

