## Supplementary Figures for "Variability vs Phenotype: multimodal analysis of Dravet Syndrome Brain Organoids powered by Deep Learning"

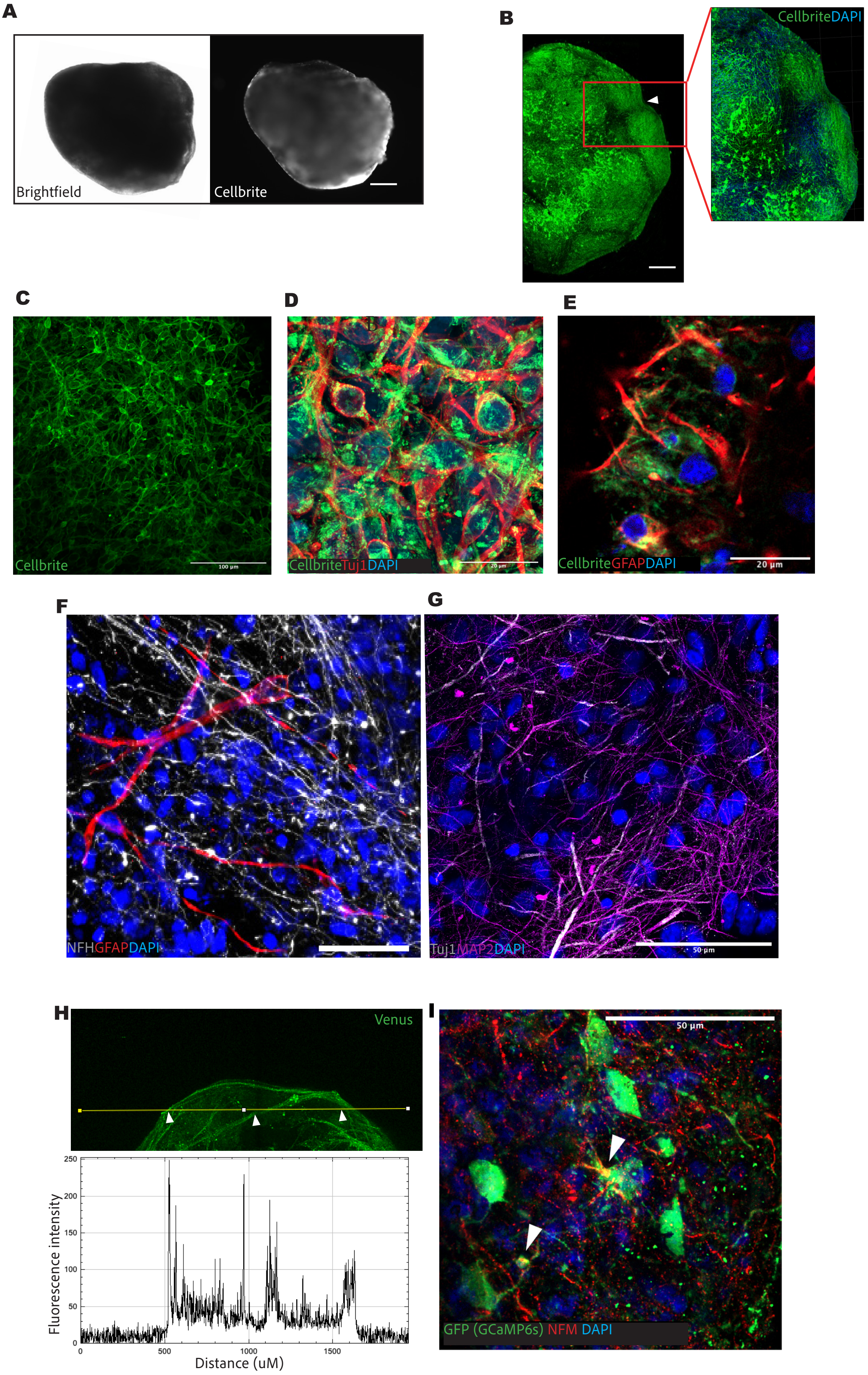

A
Brightfield
Cellbrite
B
CellbriteDAPI
C
Cellbrite
100 µm
D
CellbriteTuj1DAPI
20 µm
E
CellbriteGFAPDAPI
20 µm
F
NFHGFAPDAPI
G
Tuj1MAP2DAPI
50 µm
H
Venus
Fluorescence intensity
0
50
100
150
200
250
500
1000
1500
Distance (uM)
I
50 µm
GFP (GCaMP6s) NFM DAPI

**Supplementary Figure 1. Cellbrite staining in brain organoids.** A. Cellbrite and brightfield images of a 90-day-old H9 organoid. Scale bar: 500 µm. B. Image reconstruction of 90-day-old brain organoids stained with Cellbrite and DAPI. Scale bar: 500 µm. C. Cellbrite staining in a fixed H9 organoid. Scale bar: 100 µm. D,E. Colocalization of Cellbrite with Tuj1 (D) and GFAP (E) markers in 90-day-old brain organoids. F. Distribution of GFAP and NFM staining in 90-day-old brain organoids, revealing the spatial arrangement of astrocytes and neurons. G. Tuj1 and MAP2 staining in 90-day-old brain organoids, showing the distribution of coexisting immature and mature neurons within the organoid. H. Fluorescent signal profile of a Venus-LV-infected brain organoid at day 90 of differentiation, showing peaks of higher signal in bundle-like zones. This profile corresponds to organoid in Supplementary Video 1B. I. IF of Syn1-GCaMP6s-expressing cells together with NFM neuronal marker. GFP antibody is used to detect the GCaMP6s. Scale bar: 50 µm.

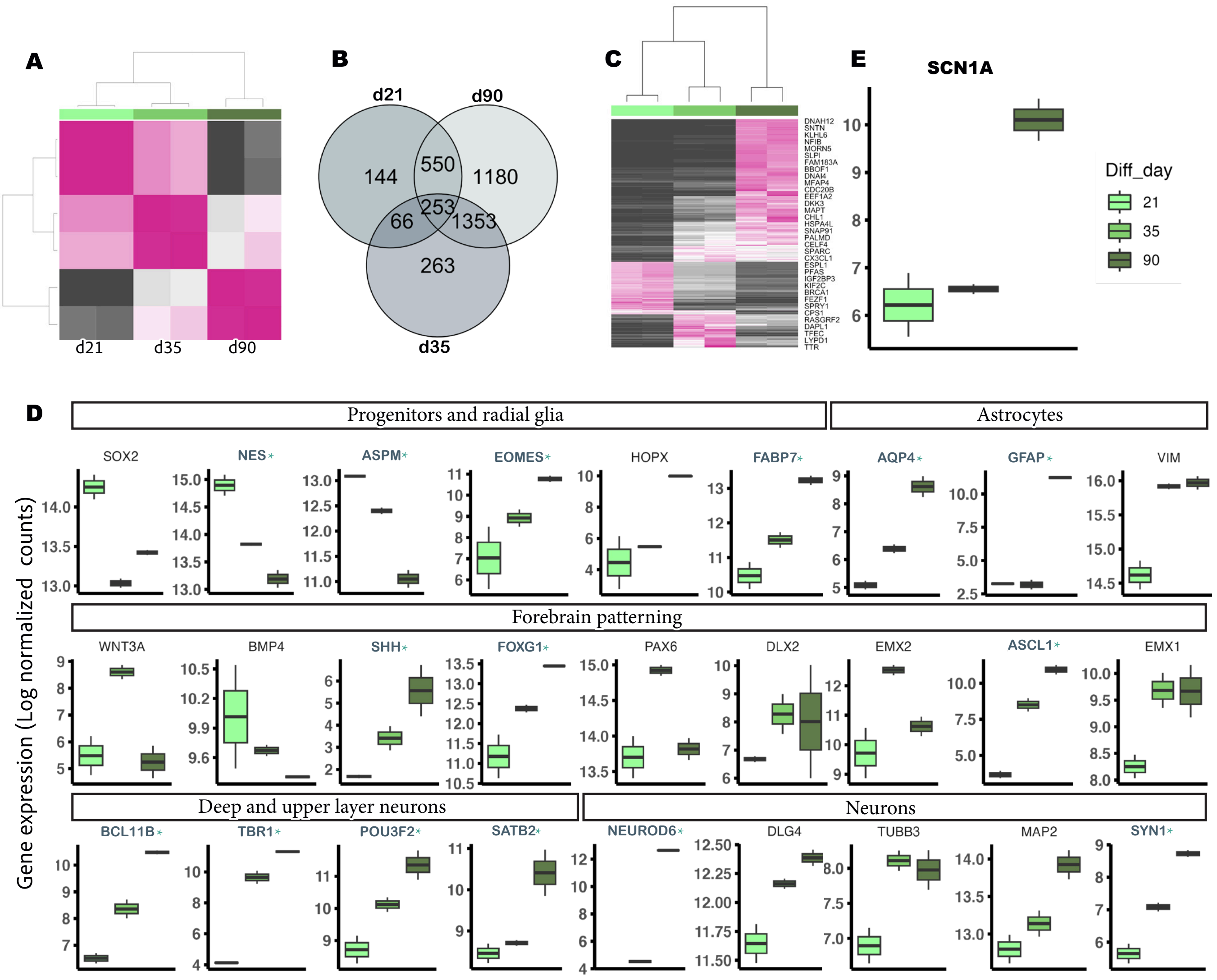


**Supplementary Figure 2. Temporal transcriptomic characterization of CTL brain organoids.** A. Transcriptomic profile distances calculated with Pearson Correlation Coefficients of day 21, 35 and 90 CTL brain organoids. B. Venn diagram including DE genes shared between day 21, 35 and 90 of differentiation in CTL brain organoids. C. Heatmap including top 150 DE genes at day 21, 35 and 90, each of them compared with the other two. The biggest differences are between day 21 and 90, however, there is a shift in the similar gene expression levels between day 21 and day 35, as well as between day 35 and day 90. D. Gene expression levels in log normalized counts of CTL brain organoids at day 21, 35 and 90 of differentiation. Typical markers for brain development are shown. Significantly expressed genes amongst the three time points are highlighted with an * (NES, ASPM, EOMES, FABP7, AQP4 , GFAP, SHH, FOXG1, ASCL1, BCL11B, TBR1, POU3F2, SATB2, NEUROD6, SYN1) e. Gene expression of SCN1A gene throughout brain organoid differentiation (day 21, 35 and 90).

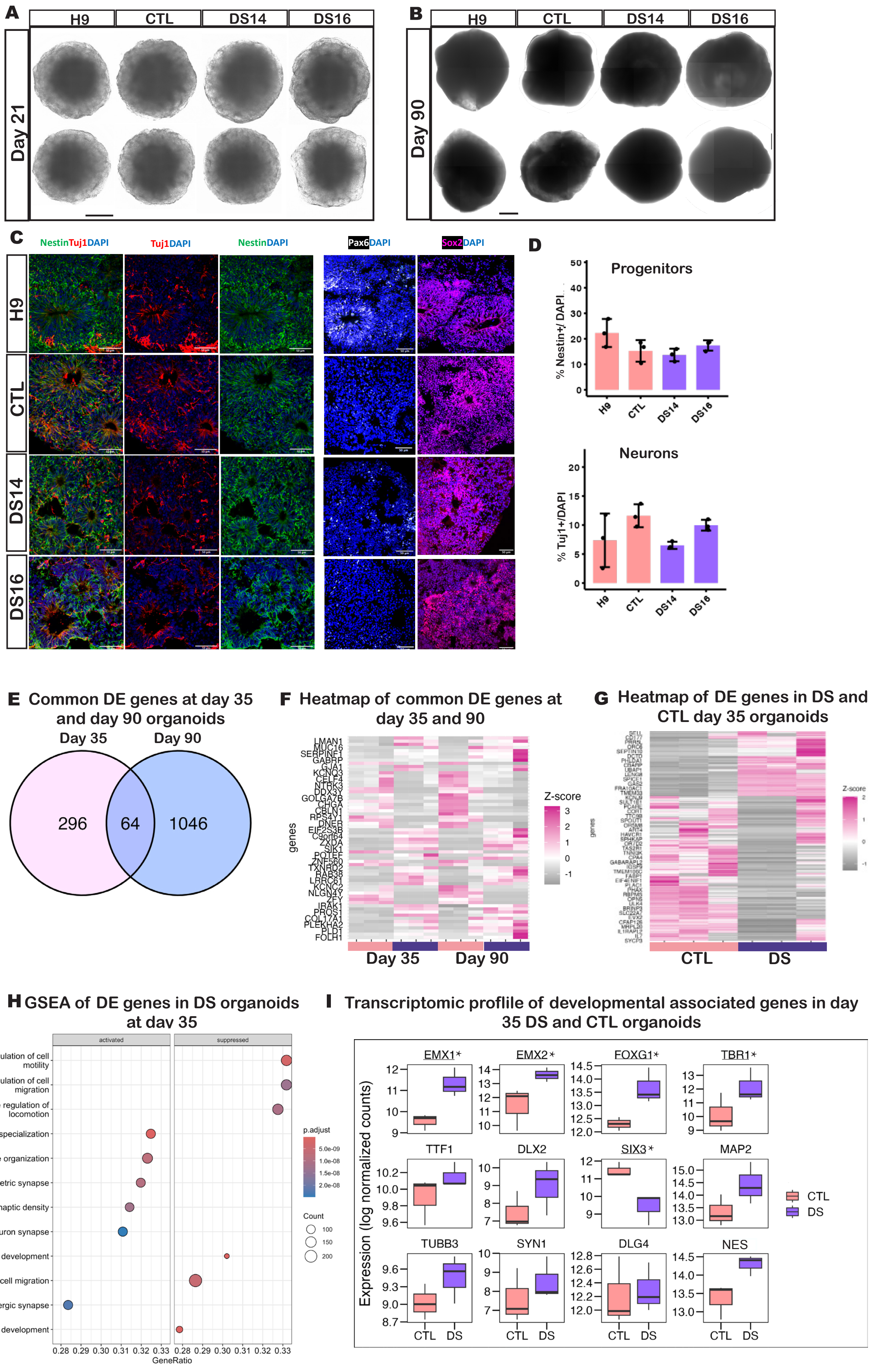
A
H9
CTL
DS14
DS16
Day 21
B
H9
CTL
DS14
DS16
Day 90
C
NestinTuj1DAPI
Tuj1DAPI
NestinDAPI
Pax6DAPI
Sox2DAPI
H9
CTL
DS14
DS16
D
Progenitors
% Nestin+/ DAPI
Neurons
% Tuj1+/DAPI
H9
CTL
DS14
DS16
E Common DE genes at day 35 and day 90 organoids
Day 35
Day 90
296
64
1046
F Heatmap of common DE genes at day 35 and 90
genes
Z-score
Day 35
Day 90
G Heatmap of DE genes in DS and CTL day 35 organoids
CTL
DS
H GSEA of DE genes in DS organoids at day 35
activated
suppressed
positive regulation of cell motility
positive regulation of cell migration
positive regulation of locomotion
postsynaptic specialization
synapse organization
asymmetric synapse
postsynaptic density
neuron to neuron synapse
skin development
regulation of cell migration
glutamatergic synapse
epidermis development
p.adjust
Count
GeneRatio
I Transcriptomic proflile of developmental associated genes in day 35 DS and CTL organoids
Expression (log normalized counts)
EMX1*
EMX2*
FOXG1*
TBR1*
TTF1
DLX2
SIX3*
MAP2
TUBB3
SYN1
DLG4
NES
CTL
DS

**Supplementary Figure 3. Complementary phenotypic characterization of brain organoids.** A,B. Brightfield images at 5x magnification of young 21 DIV (A) and mature 90 DIV (B) brain organoids. Morphological inter-organoid variability non-genotype related is observed at both time points but increased in day 90 brain organoids. Scale bar: 500um. C. IF characterization of 35-day-old H9, CTL, DS14 and DS16 brain organoids for neural progenitor cell (NPC) markers Nestin, Tuj1, Sox2 and PAX6. D. Percentage of progenitors (Nestin+) in 35-day-old brain organoids from H9, CTL, DS14 and DS16 cell lines. ANOVA test showed no statistically significant differences among cell lines (p value > 0.05). E. Percentage of immature neurons (Tuj1+) in 35-day-old brain organoids from H9, CTL, DS14 and DS16 cell lines. ANOVA test showed no statistically significant differences among cell lines (p value > 0.05). F. Venn diagram showing DE genes at day 35 that are also dysregulated at day 90. G. Heatmap of DE genes at day 90 with respect to day 35 in both DS14 and CTL brain organoids. The tendency is maintained throughout differentiation. H. Heatmap of DE genes in DS14 with respect to CTL at day 35. I. Gene set enrichment analysis (GSEA) showing upregulated (left) and downregulated (right) GO categories in DS14 with respect to CTL brain organoids both at day 35 of differentiation. Bonferroni or Benjamini-Hochberg (BH) correction was applied for calculating p adjusted value. J. Gene expression levels in log normalized counts of DS24 and CTL brain organoids at day 35 of differentiation. Typical markers for brain development are shown. Significantly expressed genes are highlighted with an * (EMX1, EMX2, FOXG1, TBR1, SIX3).

**A**

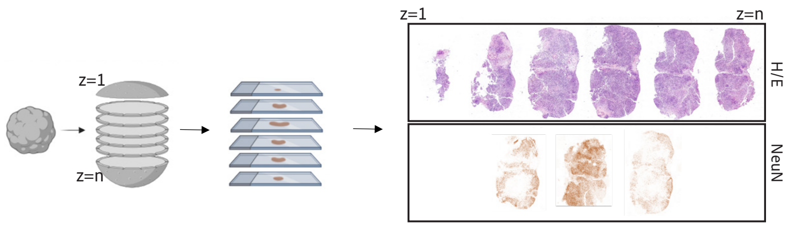


**B**

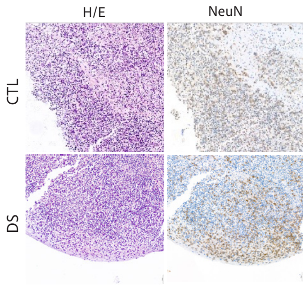


**C**

| Evaluation metric | |
|---|---|
| Accuracy | 0,732 |
| Precision | 0,733 |
| Recall | 0,735 |
| F1-score | 0,731 |

| | DRAVET | HEALTHY |
|---|---|---|
| DRAVET | 172 (0,76) | 53 (0,24) |
| HEALTHY | 80 (0,30) | 191 (0,70) |

**D**

| Evaluation metric | |
|---|---|
| Accuracy | 0,967 |
| Precision | 0,950 |
| Recall | 0,976 |
| F1-score | 0,961 |

| | DRAVET | HEALTHY |
|---|---|---|
| DRAVET | 20 (0,95) | 1 (0,05) |
| HEALTHY | 0 (0,00) | 9 (1,00) |

**E**

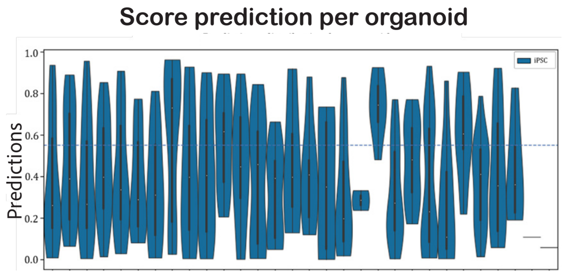


**F**

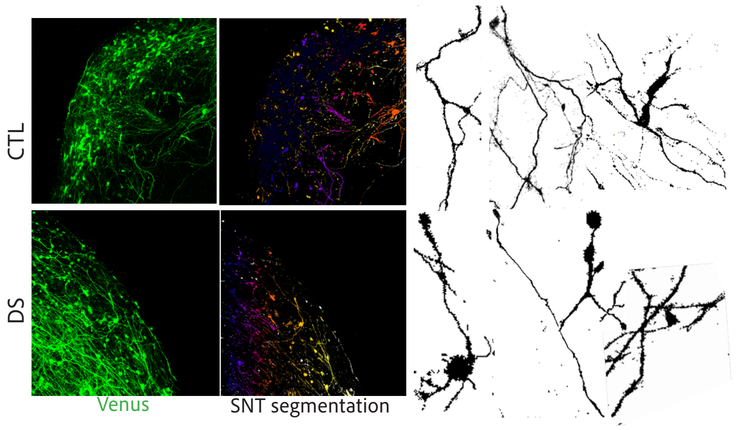


**G**

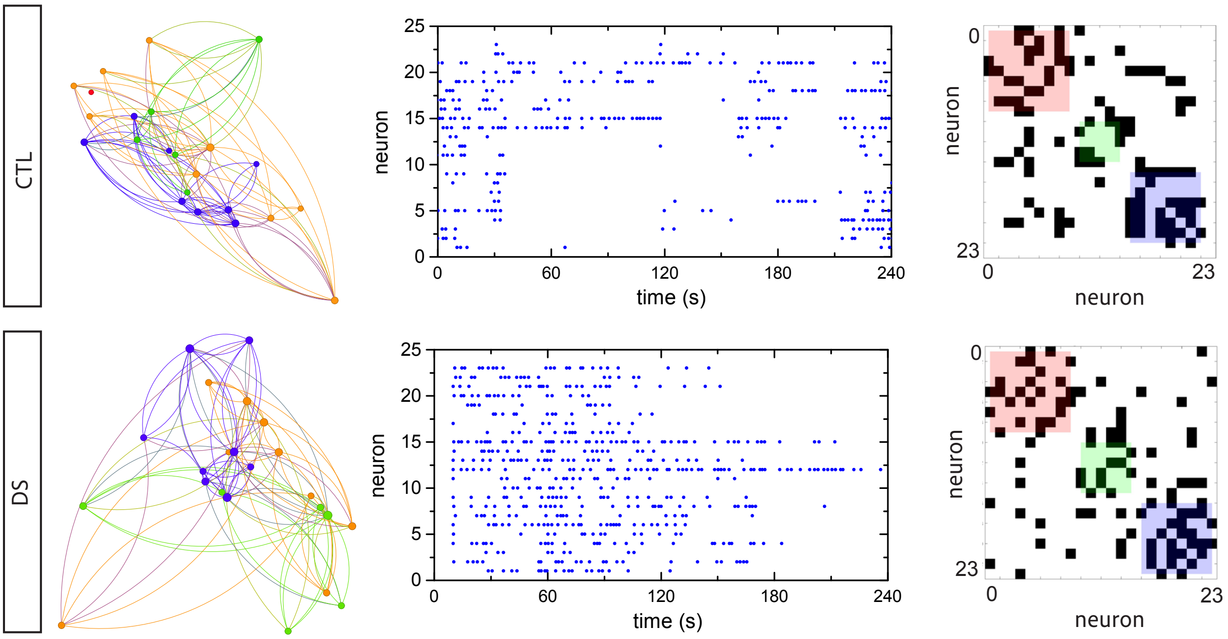

**Supplementary Figure 4. ImPheNet resolves regional and intraindividual organoid variability**. A. H/E staining and NeuN IHC of separated paraffin-sections showing variability in the composition of brain organoids along the volume. B. H/E and NeuN immunohistochemistry showing section-dependent heterogenicity of neuronal distribution. C,D. Evaluation metrics of the predictive AI-model based on fragmented (C) and whole organoid (D) scores. E. Score phenotypic predictions for each individual H9 brain organoid. The heterogenicity of the phenotypic prediction for each fragment belonging to the same organoid. F. Venus-LV infection of CTL and DS organoids showing morphology heterogenicity in individual neurons in both genotypes. G. Connectivity plots (left), raster plots (center), and connectivity matrices (right) of calcium analysis with GCaMP6s in H9 (up) and DS16 (down) brain organoids.

**A**

| Evaluation metric | |
|---|---|
| Accuracy | 0,759 |
| Precision | 0,671 |
| Recall | 0,681 |
| F1-score | 0,675 |

| | DRAVET | INHIBITORY |
|---|---|---|
| DRAVET | 242 (0,83) | 50 (0,17) |
| INHIBITORY | 42 (0,47) | 48 (0,53) |

**B**

| Evaluation metric | |
|---|---|
| Accuracy | 0,941 |
| Precision | 0,964 |
| Recall | 0,875 |
| F1-score | 0,910 |

| | DRAVET | INHIBITORY |
|---|---|---|
| DRAVET | 13 (1,00) | 0 (0,00) |
| INHIBITORY | 1 (0,25) | 3 (0,75) |

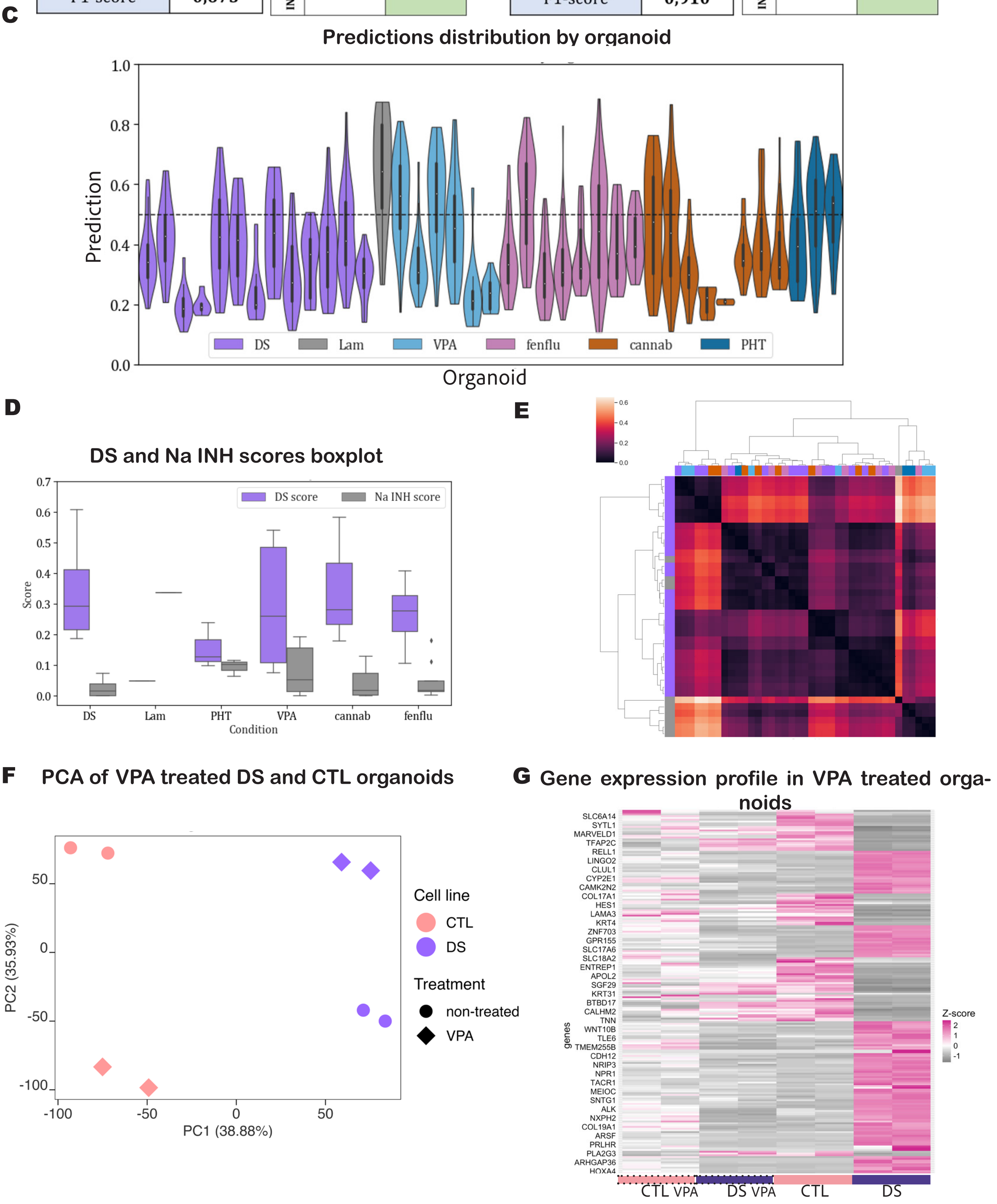

**Supplementary Figure 5. ImPheNet resolves drug effects in DS treated organoids.** A,B. Respectively, model evaluation metrics of the predictive DL-model based on A) fragmented images and B) whole organoid scores in treated organoids. C. Histogram of phenotypic predictions for each individual treated brain organoid, showing heterogenicity in the phenotypic prediction for each fragment belonging to a same organoid and suggesting a threshold for prediction of non-treated and treated organoids. D. Boxplot of the DS and INH score ratios (yellow and grey respectively) for each treatment. E. Clustermap using the Euclidean distance between the computed scores. This heatmap shows how the model scoring is clustering treatments, suggesting similarity in the treatment effects on the DS organoids. F. PCA on transcriptomic profiles of CTL and DS 90-day brain organoids untreated and treated with VPA, showing evident effect of VPA in gene expression both in CTL and in DS organoids. G Heatmap of the DE genes in DS organoids after VPA treatment that were already dysregulated in control conditions comparing DS and CTL organoids. DS organoids reach CTL organoids gene expression trend when are treated with VPA.
